# Revisiting the identification of breast cancer tumour suppressor genes defined by copy number loss of the long arm of chromosome 16

**DOI:** 10.1101/2021.07.30.454550

**Authors:** David F Callen

## Abstract

In breast cancer loss of the long-arm of chromosome 16 is frequently observed, suggesting this is the location of tumour suppressor gene or genes. Previous studies localised two or three minimal regions for the LOH genes in the vicinity of 16q22.1 and 16q24.3, however the identification of the relevant tumour suppressor genes has proved elusive. The current availability of large datasets from breast cancers, that include both gene expression and gene dosage of the majority of genes on the long-arm of chromosome 16 (16q), provides the opportunity to revisit the identification of the critical tumour suppressor genes in this region.

Utilising such data it was found 37% of breast cancers are single copy for all genes on 16q and this was more frequent in the luminal A and B subtypes. Since luminal breast cancers are associated with a superior prognosis this is consistent with previous data associating loss of 16q with breast cancers of better survival. Previous chromosomal studies found a karyotype with a der t(1;16) to be the basis for a proportion of breast cancers with loss of 16q. Use of data indicating the dosage of genes 21.9% of breast cancers were consistent with a der t(1;16) as the basis for loss of 16q. In such cases there is both loss of one dose of 16q and three doses of 1q suggesting a tumour suppressor function associated with long-arm of chromosome 16 and an oncogene function for 1q.

Previous studies have approached the identification of tumour suppressor genes on 16q by utilising breast cancers with partial loss of 16q with the assumption regions demonstrating the highest frequency of loss of heterozygosity pinpoint the location of tumour suppressor genes. Sixty one of 816 breast cancers in this study showed partial loss of 16q defined by dosage of 357 genes. There was no compelling evidence for “hot-spots” of localised LOH which would pinpoint major tumour suppressor genes. Comparison of gene expression data between various groups of breast cancers based on 16q dosage was used to identify possible tumour suppressor genes. Combining these comparisons, together with known gene functional data, allowed the identification of eleven potential tumour suppressor genes spread along 16q. It is proposed that breast cancers with a single copy of 16q results in the simultaneous reduction of expression of several tumour suppressor genes. The existence of multiple tumour suppressor genes on 16q would severely limit any attempt to pinpoint tumour suppressor genes locations based on localised hot-spots of loss of heterozygosity.

Interestingly, the majority of the identified tumour suppressor genes are involved in the modulation of wild-type p53 function. This role is supported by the finding that 80.5% of breast cancers with 16q loss have wild-type p53. TP53 is the most common mutated gene in cancer. In cancers with wild-type p53 would require other strategies to circumvent the key tumour suppressor role of p53. In breast cancers with complete loss of one dose of 16q it is suggested this provides a mechanism that contributes to the amelioration of p53 function.

## Introduction

Tumour suppressor genes (TSGs) were first proposed by Knudson (1) from his studies of retinoblastoma where individuals with a congenital mutation for one of the *RB1* alleles was at high risk of developing retinoblastoma due to a second somatic mutation of the remaining wild type *RB1* allele. Thus, the dominant wild type gene maintains normal function with inactivation of both alleles required to develop this specific cancer. Subsequently, the *TP53* gene provided another example of a TSG with individuals with a congenital mutation of one *TP53* allele being at high risk of developing a variety of solid cancers, including breast cancer (2). Analysis of these cancers showed there was somatic inactivation of the remaining wild-type allele, resulting in loss of normal function of the p53 protein in cancer cells. Subsequently, analysis of cancers, for example breast cancer (3), showed that the wild-type function of the p53 protein was frequently inactivated by mutation of both wild-type TP53 alleles, or more commonly by a combination of both mutation of one wild-type allele and loss (loss-of-heterozygosity, LOH) of the remaining wild-type allele by chromosomal mechanisms that include deletion or mitotic recombination.

On the basis of these findings, in the early 1990s the assumption seemed reasonable that if regions of the genome in breast cancers where found to have frequent LOH then this was likely to indicate that a TSG was in the vicinity (4). Since in regions of the chromosome showing frequent LOH there was varying extents of LOH in different tumours, it was considered that the location of a TSG could be defined by detailed physical mapping of these regions of LOH. The location of the TSG would be defined by the defining the minimal common region of LOH, that is the TSG would be located at the region of most frequent loss. In breast cancer early cytogenetic analysis showed loss of the long-arm of chromosome 16 (16q) was frequently observed, suggesting this is the location of a TSG (5). At this time highly polymorphic microsatellite markers were being developed and therefore use of a battery of microsatellite markers physically located on 16q could then define the extent of LOH to define the regions possible location of a TSG (6).

Use of microsatellite markers in tumour samples was technically demanding since contamination could occur from non-malignant tissue in the breast cancer samples resulting in difficulty in interpreting loss of bands resulting from LOH. It should be noted that this technique determines LOH that can be the result of either physical loss of the region of the chromosome (resulting in a haploid region) or that due to mitotic recombination which can generate regions of the chromosome where both homologues are identical and therefore homozygous. These are often referred to as copy number loss and copy-neutral respectively. Some indication of the frequency of these two mechanisms can be gauged in a small study of 20 breast tumours where 25% of cases of 16q LOH were due to mitotic recombination (7).

The first detailed cytogenetic characterisations of breast cancers identified the presence of multiple cases with a der(1;16)(q10;p10) in a karyotype with two intact chromosome 1s and a single copy of chromosome 16. Overall this results in trisomy for 1q and monosomy for 16q (5). FISH characterisation of such der(1;16)s using satellite probes from centromeric regions showed the rearrangements occurred in various regions of the centromeric repeat regions of chromosomes 1 and 16 (8). There are a number of candidate and known oncogenes on 1q that could contribute to breast oncogenesis where elevated expression is associated with 1q trisomy (9). While loss of 16q in breast tumours was most frequently associated with low tumour grade and estrogen receptor (ER) positivity (luminal breast cancers) defining TSG proved elusive (10). Series of breast cancers were analysed with a panels of microsatellite markers on 16q (11,12) with the largest study including 712 breast cancers (13). These studies localised two or three minimal regions for the TSGs in the vicinity of 16q22.1 and 16q24.3. However, results were not unambiguous and the rare cases of very localised 16q LOH may also include regions of loss due to small deletions of complex rearrangements unrelated to the presence of TSGs.

The current availability of large datasets from breast cancers that include both gene expression and gene dosage of the majority of 16q genes provides the opportunity to revisit the LOH of 16q and determine if this now provides data to allow the identification of the critical TSGs located in this region. This study will also investigate the reported associated of LOH of 16q with a favourable prognosis and luminal breast cancers and if copy number loss of 16q in breast cancer is often the consequence of a karyotype with a derivative t(1;16) that results in trisomy for 1q and monosomy for 16q.

## Materials and Methods

The analysis was based on the TCGA data set of molecular profiles of 816 breast cancer cases (14). A list of all genes on the long arm of chromosome 16 (16q) was downloaded from the NCBI data base. Forty-one genes with absent or incomplete data in the TCGA dataset were eliminated from the analysis. All microRNAs, SNORDs and other non-coding RNAs, pseudogenes and other unverified coding genes were also eliminated resulting in a final list of 357 genes located on 16q. A similar approach was used to determine dosage of chromosome 1q and 16p in the investigation of the presence of a possible der(1;16) translocation.

A subset of the breast cancers were classified by the PAM-50 criteria into Luminal A, Luminal B, HER2+ positive and Basal subgroups (15) and this classification was used for additional analyses. Genomic Identification of Significant Targets in Cancer (GISTIC) and RNA sequence (RNAseq V2 RSEM) were downloaded for the breast cancers analysed. LOH of the long arm of chromosome 16 was defined when the 357 16q genes analysed had a GISTIC value of -1, indicating a single dose of 16q and were termed “entire 16q loss”. Where all 357 genes had a GISTIC value of zero this indicated two intact doses of 16q and were termed “diploid for 16q”. Where GISTIC values of all genes were either -1 or 0 the breast cancers were classified as “partial 16q loss”. The remaining breast cancers were characterised by complex dosages of 16q genes and were termed “complex 16q dosage”. Gene expression was analysed using RNA sequence data (RSEM).

## Results and Discussion

### Assessing dosage of chromosome 16 from molecular data

The dosage of the long arm of chromosome 16 in 816 breast cancer cases (14) was assessed from Genomic Identification of Significant Targets in Cancer (GISTIC) values of 357 genes (16), Table 1. Diploid 16p or 16q was defined by all 357 genes for 16q, or 413 genes for 16p, having a GISTIC value of 0, and entire 16p or 16q loss by a GISTIC value of -1 for all genes on the respective chromosomal arm. Partial 16 p or q loss was identified when all genes on the respective arm had GISTIC values of -1 or 0. Triploid dosage was defined by a GISTIC value of 1 for all genes on the respective 16 chromosomal arms. The remaining breast cancers were termed complex dosage for that arm.

**Table 1.** Dosage changes of chromosome 16 in breast cancer

| Dosage of 16 arm | 16p: Number of breast tumours | 16q: Number of breast tumours |
| --- | --- | --- |
| Entire loss | 32 (3.9%) | 302 (37.0%) |
| Diploid | 233 (28.6%) | 147 (18.0%) |
| Triploid | 322 (39.5%) | 0 |
| Partial loss | 24 (2.9%) | 61 (7.5%) |
| Complex dosage | 175 (25.1%) | 306 (37.5%) |
| Total | 816 | 816 |

Similarly, GISTIC values of 413 genes on 16p were analysed to assess the dosage of 16p. Interestingly there was a group of five genes, *TP53TG3B, TP53TG3C, TP53TG3, TP53TG3D* and *ZNF267* adjacent to the centromere in a region of approximately 1.5 Mb that was apparently deleted in 23.8% of breast cancers. In 63 cases these five genes were in a single dose while all other 16p genes were two doses (consistent with diploid 16p) and in 98 cases the five genes were in two doses while all other 16p genes had three doses (consistent with triploid 16p). The mRNA data was consistent with an absence of expression of the *TP53TG3* cluster in breast tumours. There was expression of *ZNF267* which has been reported to be upregulated in hepatocellular carcinoma (17). However, in breast cancers there is no apparent upregulation of *ZNF267* as expression was not related to whether there was a one or two doses of *ZNF267* since average mRNA expression in breast cancers with a single dose if 16p was 388.0 and in cancers with two doses of 16p 398.7 (p=0.46, not significant). However, from the database of genetic variants (18) this region is reported to be associated with frequent deletions/duplications in normal populations. It has previously been reported that human copy number variants are significantly overrepresented close to telomeres and centromeres and in simple tandem repeat sequences (19). It is considered that this region is more likely to represent a common copy variant in the normal population rather than any breast cancer specific event.

There were 18% of the 816 breast cancers that were diploid (GISTIC=0) for all the 357 16q genes analysed while 37.5% of breast cancers had complex regions of 16q gene dosage with regions of 16q ranging from occasional apparent complete loss to partial triploid. In 7.5% of tumours there was partial “simple” loss of 16q, defined as tumours where all genes on 16q exhibited either diploid or single copy dosages. In these cancers the physical regions involved in 16q loss ranged from contiguous to non-contiguous regions and ranged in size from single gene loss to loss of the majority of 16q. In contrast, 16p was rarely haploid (3.9%) but frequently triploid (39.5%). In the 37% of tumours that were single copy for all 16q genes 50% had three doses for 16p, 21% two doses and 7% a single dose.

A subgroup of the breast tumours were classified by the Pam-50 criteria and it was determined if the various categories of 16q loss varied between these subgroups, Table 2. The proportion of breast cancers with 16q loss of the entire long arm was not significantly different for all four subtypes of breast cancers. However, the pooled luminal A and B 16q loss (39.4%) was significantly 2.5-fold greater (p<0.0001) than the pooled basal-like and HER2 enriched 16q loss (15.8%). Rye et al (20) have used quantitative FISH of breast carcinomas with known molecular subtypes to determine the loss of 16q. These studies showed whole arm loss of 16q occurred in 40% of luminal A tumours but only 22% of basal-like tumours. These data are highly comparable with the data presented in Table 2 estimated from the GISTIC data and provide an independent validation of the relevance of the GISTIC data.

**Table 2.** Dosage changes of 16q in the PAM-50 subtypes of breast cancer

| Dosage of 16q | PAM-50 classification (number of cases and %) |  |  |  | Total |
| --- | --- | --- | --- | --- | --- |
|  | Luminal A | Luminal B | Basal-like | HER2 enriched |  |
| Diploid | 33 (16.5%) | 17 (13.9%) | 13 (12.2%) | 12 (23.5%) | 75 (15.6%) |
| Entire loss | 80 (40.0%) | 47 (38.5%) | 13 (12.2%) | 11 (21.6%) | 151 (31.5%) |
| Partial loss | 16 (8.0%) | 9 (7.4%) | 12 (11.2%) | 3 (5.9%) | 40 (8.3%) |
| Complex dosage | 71 (35.5%) | 49 (40.2%) | 69 (64.5%) | 25 (49.0%) | 214 (44.6%) |
| Total | 200 | 122 | 107 | 51 |  |

### Derivative t(1;16)

Cytogenetic investigations of breast cancers were the first to identify the presence of a derivative chromosome resulting from a translocation with breakpoints within the peri-centromeric heterochromatin of chromosomes 1 and 16, a der t(1;16). This appears to be an unbalanced translocation as only the derivative chromosome consisting of a fusion between 1q and 16p is found. This der t(1;16) is typically in a karyotype with two normal chromosome 1s and a single chromosome 16 resulting in overall trisomy for 1q and monosomy for 16q (5) (8).

The GISTIC data was evaluated to determine if a der t(1;16) is likely to be present in the breast cancer cases. Using GISTIC data for genes on the long arm of chromosome 1 there were 112 cases (13.7%) where genes on 1q were present in three doses and in addition dosage of 1p genes (genes from 1p33 to the centromere were assessed) were in two doses. Interestingly, three doses of 1q occurred in 21.9% of breast cancers with loss of the entire 16q while 1q trisomy was more than two-fold less in the other two categories and absent in cases with partial loss of chromosome 16q (Table 3). Within the luminal A subgroup the frequency of an apparent der t(1;16) was 23% while in the basal-like subgroup only 3.7%. This is comparable with the data from chromosomal fluorescence *in situ* hybridisation studies where 27% of luminal A but only 2% basal-like tumours possessed the der t(1;16) (20).

Previous studies indicated the der t(1;16) was typically present in a karyotype with a single chromosome 16. GISTIC values for 16p genes demonstrated that 25.8% of the 66 cases with the presumptive der t(1;16) had two doses of 16p while 66.7% of cases there were three doses. A simplistic scenario for the karyotype of breast cancers with the der t(1;16) as the basis for 16q LOH would be two normal chromosome 1s, a normal chromosome 16 and the der t(1;16) consisting of 16p and 1q. However, breast cancer karyotypes are generally complex and detailed determination of chromosome rearrangements from GISTIC data is not possible.

**Table 3.**
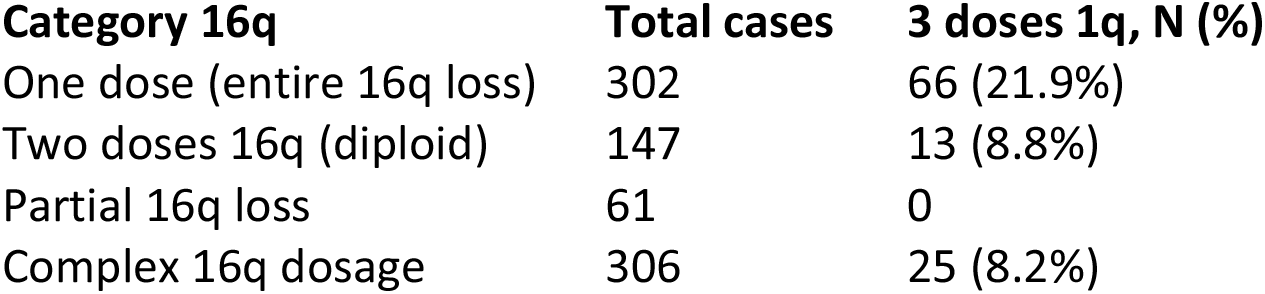
Frequency of possible der t(1;16) as assessed from GISTIC values

### Are there “hotspots” of LOH on 16q?

In previous studies, following from the observation of frequent LOH of 16q in breast cancers extensive efforts were undertaken to map regions of LOH to determine a “minimum region of overlap” between LOH regions that would locate the possible location of a TSG (13). This was based on the premise that the observation of LOH indicated the presence of a TSG and its location could be pin-pointed by more detailed analysis of breast cancer cases with partial LOH. An implied assumption was that one copy of the TSG would be lost through LOH while the other copy would be inactivated by mutation or methylation. Such studies, using classical polymorphic microsatellite markers, suggested two possible locations for TSGs, 16q22.1 and 16q24.3, as these were regions with the highest frequency of localised LOH.

In the cohort of 816 breast cancers, the availability of GISTIC data from 357 genes on 16q provided enhanced resolution to map regions of dosage loss compared with any previous studies. There were 61 breast cancers where one or more 16q genes were consistent with haploid dosage (single copy gene loss) and where all other 16q genes were diploid (present with two copies). These were termed partial 16q loss. The regions varied from single gene loss to loss of the majority of 16q. Using the genomic location of each gene, the frequency of gene loss among the subgroup of 61 breast cancers with partial 16q loss was plotted across the entire length of the long-arm, Figure 1.

**Fig 1.**
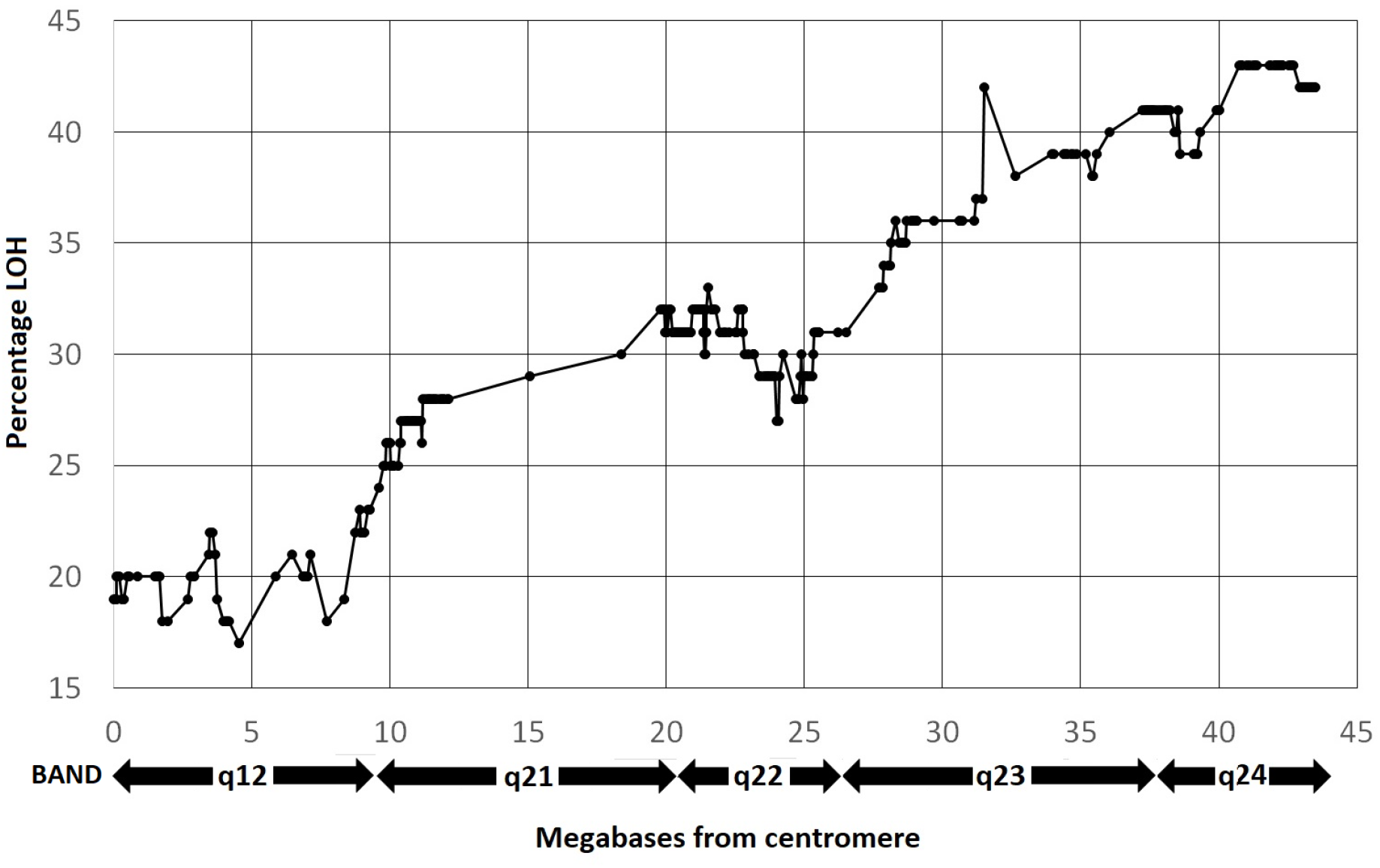
Frequency of gene single copy loss among breast cancer cases with partial 16q loss.

The genes are plotted on the X axis in relation to their physical position (megabases) from centromere to 16qter (left to right). The approximate locations of chromosome bands are indicated on the X axis.

The data does not support regions of peak loss at 16q22.2 and 16q24.3 as previously reported but rather a loss that gradually increases from the centromere to the telomere. If breakage of 16q is a random event then the increase would be expected to be linear. Regression analysis showed the data deviated significantly from a linear increase. However, breakage events are likely to be influenced by local areas of DNA sequence with inherent fragility (21) resulting in deviations from a linear trajectory. There is no compelling evidence for “hot-spots” of localised LOH which would pinpoint a major TSG. A possible exception is a peak at 31.52 megabases which corresponds to the location of the *WWOX* gene, previously reported as a known tumour suppressor gene in breast cancer (22). Of the 61 partial LOH cases in five of these the loss involved a single gene with three of these being *WWOX* and with one case involving loss of *SALL1* and one of *WDR59*. There was a further case with loss of just two genes, *WWOX* and *FAM32B* which are located approximately 7 Mb apart. Loss of *WWOX* in other breast cancers was associated with large regions of loss.

Since mutations of *WWOX* in breast cancer have not been reported, if *WWOX* was a significant TSG in breast cancer this gene would be predicted to demonstrate haplo-insufficiency. *WWOX* has been shown to drive metastasis in triple-negative breast cancers (23) and indeed two of the four cancers with loss restricted to *WWOX* were triple-negative cancers. For the breast cancers with loss of the entire 16q, which are predominately of the luminal subtype, expression of *WWOX* is not significantly altered by one or two doses of the gene (average expression 389.7 RMEM for cases with two doses of 16q genes vs average 326.6 for single dose of all 16q genes, p=0.11, ns). *WWOX* spans the common chromosomal fragile site *FRA16D* and this localised fragility is likely to contribute to incidents of specific loss of *WWOX*.

### Relationship between copy number and gene expression

Classical tumour suppressor genes are frequently functionally compromised by loss of one allele by a physical chromosome loss with subsequent mutation of the remaining allele. However there is a class of TSG where reduction of gene dosage from two copies to a single copy is sufficient to reduce transcriptional activity and compromise the normal functioning of the gene in tumours (24). There are no highly mutated genes in breast cancer located on 16q (25) although BC driver genes, *CBFB, CDH1, CTCF* and *NUP93*, have been identified (, 26). It is suggested that loss of transcriptional activity associated with copy number loss is likely to be the basis for functionally compromising the function of TSGs in this region.

In an approach to detect potential TSGs on 16q RNAseq data was utilised as a measure of gene expression. Two approaches were used to identify potential TSGs. Firstly, it was considered that TSGs would be downregulated in luminal A and B tumours with LOH of the entire 16q long-arm compared with those tumours with diploid doses of 16q genes. This identified 175 genes significantly downregulated (p<0.05 with Boneferri correction). Secondly, since the incidence of complete loss of 16q was frequent in the luminal A and B subtypes but significantly lower in the basal-like and HER2 enriched PAM50 subtypes it was considered that 16q TSGs would unlikely to be downregulated in the latter breast cancers. Comparison of RNAseq data for all genes on 16q identified 190 genes significantly downregulated (p<0.05 with Boneferri correction) in the luminal A and B tumours with a single dose of all 16q genes compared with basal-like and HER2 enriched cancers. When these two sets of downregulated genes were compared there were 132 in common (69% of genes from the comparison with basal and HER2 subtypes, and 75% from the comparison within the luminal subtypes).

From these relative comparisons of gene expression oncogenes in addition to TSGs will also be potentially detected. For example, genes shown to be down-regulated in luminal A and B subtypes with loss of 16q may represent those with an oncogene function specifically in luminal A and B subtypes with diploid 16q. Similarly, in the comparison of basal-like and HER2 enriched cancers with luminal cancers with loss of 16q, down-regulated genes in the luminal subtypes can be the result of upregulated oncogenic functions in basal-like and HER2 enriched cancers. Therefore. for each of the 132 genes in common between the two sets of data literature searches for each gene were undertaken to determine evidence for a potential role as a TSG. This analysis allowed detection of oncogenic genes eg *GINS2* upregulated in basal-like cancers. Genes were eliminated as candidate TSGs due to lack of definitive evidence of function or relationship to cancer, absence of any evidence for a tumour suppressor role or a function more consistent with an oncogenic role. Eleven genes were considered to have evidence supporting a TSG role and these genes were not clustered but distributed along the length of 16q (Table 4).

**Table 4.**
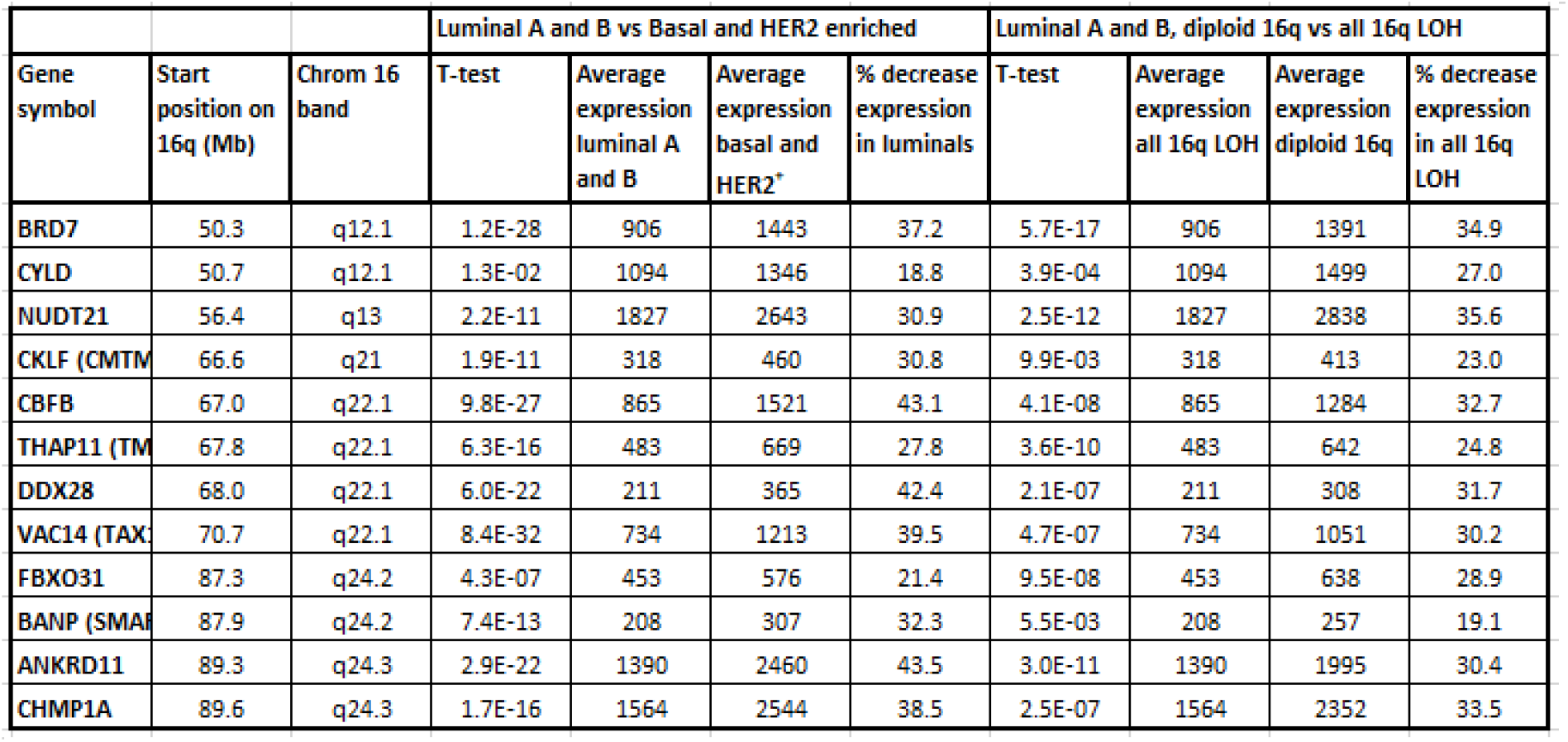
Potential tumour suppressor genes on the long arm of chromosome 16.

| Gene symbol | Start position on 16q (Mb) | Chrom 16 band | Luminal A and B vs Basal and HER2 enriched |  |  |  | Luminal A and B, diploid 16q vs all 16q LOH |  |  |  |
| --- | --- | --- | --- | --- | --- | --- | --- | --- | --- | --- |
|  |  |  | T-test | Average expression luminal A and B | Average expression basal and HER2* | % decrease expression in luminals | T-test | Average expression all 16q LOH | Average expression diploid 16q | % decrease expression in all 16q LOH |
| BRD7 | 50.3 | q12.1 | 1.2E-28 | 906 | 1443 | 37.2 | 5.7E-17 | 906 | 1391 | 34.9 |
| CYLD | 50.7 | q12.1 | 1.3E-02 | 1094 | 1346 | 18.8 | 3.9E-04 | 1094 | 1499 | 27.0 |
| NUDT21 | 56.4 | q13 | 2.2E-11 | 1827 | 2643 | 30.9 | 2.5E-12 | 1827 | 2838 | 35.6 |
| CKLF (CMTM) | 66.6 | q21 | 1.9E-11 | 318 | 460 | 30.8 | 9.9E-03 | 318 | 413 | 23.0 |
| CBFB | 67.0 | q22.1 | 9.8E-27 | 865 | 1521 | 43.1 | 4.1E-08 | 865 | 1284 | 32.7 |
| THAP11 (TM) | 67.8 | q22.1 | 6.3E-16 | 483 | 669 | 27.8 | 3.6E-10 | 483 | 642 | 24.8 |
| DDX28 | 68.0 | q22.1 | 6.0E-22 | 211 | 365 | 42.4 | 2.1E-07 | 211 | 308 | 31.7 |
| VAC14 (TAX) | 70.7 | q22.1 | 8.4E-32 | 734 | 1213 | 39.5 | 4.7E-07 | 734 | 1051 | 30.2 |
| FBXO31 | 87.3 | q24.2 | 4.3E-07 | 453 | 576 | 21.4 | 9.5E-08 | 453 | 638 | 28.9 |
| BANP (SMA) | 87.9 | q24.2 | 7.4E-13 | 208 | 307 | 32.3 | 5.5E-03 | 208 | 257 | 19.1 |
| ANKRD11 | 89.3 | q24.3 | 2.9E-22 | 1390 | 2460 | 43.5 | 3.0E-11 | 1390 | 1995 | 30.4 |
| CHMP1A | 89.6 | q24.3 | 1.7E-16 | 1564 | 2544 | 38.5 | 2.5E-07 | 1564 | 2352 | 33.5 |

Evidence for TSG function is briefly presented for each of the eleven genes listed in Table 4.

- The BRD7 protein can be recruited to target gene promotors and influences promotor activity. In particular BRD7 is a p53 cofactor required for the efficient induction of p53-dependent oncogene-induced senescence (27). Evidence suggests BRD7 blocks tumour growth, migration and metastasis by negatively regulating YB1-induced EMT (28).
- CYLD has also, similar to BRD7, been implicated in p53 function as it regulates the p53 DNA damage response (29) and is involved in mammary epithelial-mesenchymal transition (EMT) (30).
- NUDT21 is downregulated in breast cancer tissues, and also has possible involvement in EMT (31).
- CBFB is a transcription factor that binds to mRNAs and enhances transcription. In particular, in breast cancer the pathway involving CBFB binding of RUNX1 mRNA is of significance since this represses the oncogenic NOTCH signalling pathway. CBFB is a mutated driver in a variety of cancers including breast cancer (32). In breast cancer cells the downregulation of CBFB results in evasion of both translation and transcription surveillance and CBFB cooperates with p53 to maintain TAp73 expression and suppress breast cancer (33).
- THAP11 functions as a TSG in various cancer cell lines, including the breast cancer cell line MCF7, as down-regulation results in growth suppression by negatively regulating the expression of c-Myc (34).
- It has been shown in a hypoxic glioblastoma cell line that DDX28 can function as a TSG by repressing HIF-2α- and eIF4E2-mediated translation activation of oncogenic mRNAs (35). Since DDX28 protein levels relative to levels in normal tissue are reduced in several cancers, including breast cancer, this gene was considered as a potential TSG in breast cancer.
- VAC14 (TAX1BP2) is a centrosomal protein and reported as a TSG in hepatocellular carcinoma. Investigations show VAC14 can upregulate p53 by activating p38 and is also stabilised by ATM phosphorylation (36). There are no studies directly relevant to breast cancer.
- FBXO31 is a cell-cycle regulated protein and has been reported as a TSG in breast cancer and other cancers. Functionally the protein contributes to its tumour suppressor function be several mechanisms including targeting the DNA replication factor CDT1 (37).
- BANP(SMAR1) is a nuclear matrix associated protein and several mechanisms have been proposed to contribute to a tumour suppressor function. These include promoting epithelial to mesenchymal transition via E-cadherin up-regulation (38) and p53 modulation (39).
- ANKRD11 has been shown to be a p53 co-activator and can suppress the oncogenic potential of mutant p53 (40).
- There is no published data relating to the role of CHMP1A in breast cancer although a TSG role has been shown in pancreatic cancer where its role Is potentially through the p53 signalling pathway (41).

Of the eleven 16q genes identified with evidence for a role as a TSG, seven have a role in the modulation of the p53 signalling pathways. On this basis it is proposed that one of the major roles for the TSGs on 16q in the normal cell is to combine to maintain the normal role of p53 in suppressing oncogenesis. The suppression of normal p53 function in a tumour with wild-type p53 is not well understood and is expected to be multifactorial. It is suggested loss of 16q, by reducing the level of transcription, compromises the function of a number of genes on 16q that are normally required to maintain p53 function. This is consistent with the finding that LOH of 16q is associated with the early development of breast cancer (5). As a consequence of this proposal it would be predicted that wild-type p53 would be more frequent in those breast cancer cases with LOH of the entire 16q compared with those breast cancers with a diploid dose of all 16q genes. This indeed was the case with wild-type p53 found in 80.5% of breast cancers with LOH of the entire 16q compared with 68.7% which are diploid for 16q (p=0.008).

## Conclusion

The GISTIC data from the TCGA gene dosage in breast cancers provides measurement of gene dosage in a large cohort of breast cancers. Utilising these data 37% of breast cancers were single copy for all 16q genes and this is more frequent in the luminal A and B subtypes. Since luminal breast cancers are associated with a superior prognosis this is consistent with previous data associating loss of 16q with breast cancers of better survival. Previous chromosomal studies found a karyotype with a der t(1;16) to be the basis for a proportion of breast cancers with loss of 16q. Use of GISTIC data for chromosomes 1 and 16 showed 21.9% of breast cancers were consistent with a der t(1;16) as the basis for loss of 16q. In such cases there is both loss of 16q and three doses of 1q suggesting a tumour suppressor function associated with 16q and an oncogene function for 1q.

Previous studies have approached the identification of TSGs on 16q by utilising breast cancers with partial loss of 16q with the assumption regions demonstrating the highest frequency of loss of heterozygosity pinpoint the location of TSGs. Sixty one of 816 breast cancers in this study showed partial loss of 16q defined by dosage of 357 genes. There was no compelling evidence for “hot-spots” of localised LOH which would pinpoint a major TSG. A possible exception was a small peak encompassing the *WWOX* gene but this was an unlikely candidate based on expression data. Since *WWOX* spans the common chromosomal fragile site *FRA16D* this localised fragility was considered to contribute to instances of specific loss of *WWOX*.

To identify possible TSGs on 16q gene expression data was compared between breast cancers with one dose of all 16q genes and those with two doses of all 16q genes. In addition, breast cancers with 16q loss were also compared with basal-like and HER2 enriched cases. Combining these comparisons, together with known gene functional data, allowed the identification of eleven potential TSGs spread along 16q. It is proposed that breast cancers with a single copy of 16q results in the simultaneous reduction of expression of several TSGs. The existence of multiple TSGs on 16q would severely limit any attempt to pinpoint TSG locations based on localised hot-spots of loss of heterozygosity.

Interestingly, the majority of the identified TSGs are involved in the modulation of wild-type p53 function. This role is supported by the finding that 80.5% of breast cancers with 16q loss have wild-type p53. TP53 is the most common mutated gene in cancer. In cancers with wild-type p53 would require other strategies to circumvent the key tumour suppressor role of p53. In breast cancers with complete loss of one dose of 16q it is suggested this provides a mechanism that contributes to the amelioration of p53 function.

## Notes

### Competing Interest Statement

The authors have declared no competing interest.

## References

1. Knudson AG, Jr. Heredity and human cancer. The American journal of pathology. 1974;77(1):77–84.

2. Easton D, Ford D, Peto J. Inherited susceptibility to breast cancer. Cancer surveys. 1993;18:95–113.

3. Callahan R, Cropp C, Sheng ZM, Merlo G, Steeg P, Liscia D, et al. Definition of regions of the human genome affected by loss of heterozygosity in primary human breast tumors. Journal of cellular biochemistry Supplement. 1993;17G:167–72.

4. Devilee P, Cornelisse CJ. Somatic genetic changes in human breast cancer. Biochimica et biophysica acta. 1994;1198(2-3):113-30.

5. Dutrillaux B, Gerbault-Seureau M, Zafrani B. Characterization of chromosomal anomalies in human breast cancer. A comparison of 30 paradiploid cases with few chromosome changes. Cancer genetics and cytogenetics. 1990;49(2):203–17.

6. Sato T, Tanigami A, Yamakawa K, Akiyama F, Kasumi F, Sakamoto G, et al. Allelotype of breast cancer: cumulative allele losses promote tumor progression in primary breast cancer. Cancer research. 1990;50(22):7184–9.

7. Cleton-Jansen AM, Buerger H, Haar N, Philippo K, van de Vijver MJ, Boecker W, et al. Different mechanisms of chromosome 16 loss of heterozygosity in well-versus poorly differentiated ductal breast cancer. Genes, chromosomes & cancer. 2004;41(2):109–16.

8. Kokalj-Vokac N, Alemeida A, Gerbault-Seureau M, Malfoy B, Dutrillaux B. Two-color FISH characterization of i(1q) and der(1;16) in human breast cancer cells. Genes, chromosomes & cancer. 1993;7(1):8–14.

9. Muthuswami M, Ramesh V, Banerjee S, Viveka Thangaraj S, Periasamy J, Bhaskar Rao D, et al. Breast tumors with elevated expression of 1q candidate genes confer poor clinical outcome and sensitivity to Ras/PI3K inhibition. PloS one. 2013;8(10):e77553.

10. Burger H, de Boer M, van Diest PJ, Korsching E. Chromosome 16q loss--a genetic key to the understanding of breast carcinogenesis. Histology and histopathology. 2013;28(3):311–20.

11. Tsuda H, Callen DF, Fukutomi T, Nakamura Y, Hirohashi S. Allele loss on chromosome 16q24.2-qter occurs frequently in breast cancers irrespectively of differences in phenotype and extent of spread. Cancer Res. 1994;54(2):513–7.

12. Cleton-Jansen AM, Moerland EW, Kuipers-Dijkshoorn NJ, Callen DF, Sutherland GR, Hansen B, et al. At least two different regions are involved in allelic imbalance on chromosome arm 16q in breast cancer. Genes, chromosomes & cancer. 1994;9(2):101–7.

13. Cleton-Jansen AM, Callen DF, Seshadri R, Goldup S, McCallum B, Crawford J, et al. Loss of heterozygosity mapping at chromosome arm 16q in 712 breast tumors reveals factors that influence delineation of candidate regions. Cancer Res. 2001;61(3):1171–7.

14. Ciriello G, Gatza ML, Beck AH, Wilkerson MD, Rhie SK, Pastore A, et al. Comprehensive Molecular Portraits of Invasive Lobular Breast Cancer. Cell. 2015;163(2):506–19.

15. Parker JS, Mullins M, Cheang MC, Leung S, Voduc D, Vickery T, et al. Supervised risk predictor of breast cancer based on intrinsic subtypes. J Clin Oncol. 2009;27(8):1160–7.

16. Beroukhim R, Getz G, Nghiemphu L, Barretina J, Hsueh T, Linhart D, et al. Assessing the significance of chromosomal aberrations in cancer: methodology and application to glioma. Proceedings of the National Academy of Sciences of the United States of America. 2007;104(50):20007–12.

17. Schnabl B, Valletta D, Kirovski G, Hellerbrand C. Zinc finger protein 267 is up-regulated in hepatocellular carcinoma and promotes tumor cell proliferation and migration. Experimental and molecular pathology. 2011;91(3):695–701.

18. MacDonald JR, Ziman R, Yuen RK, Feuk L, Scherer SW. The Database of Genomic Variants: a curated collection of structural variation in the human genome. Nucleic Acids Res. 2014;42(Database issue):D986–92.

19. Nguyen DQ, Webber C, Ponting CP. Bias of selection on human copy-number variants. PLoS genetics. 2006;2(2):e20.

20. Rye IH, Lundin P, Maner S, Fjelldal R, Naume B, Wigler M, et al. Quantitative multigene FISH on breast carcinomas identifies der(1;16)(q10;p10) as an early event in luminal A tumors. Genes, chromosomes & cancer. 2015;54(4):235–48.

21. Bignell GR, Greenman CD, Davies H, Butler AP, Edkins S, Andrews JM, et al. Signatures of mutation and selection in the cancer genome. Nature. 2010;463(7283):893–8.

22. Pospiech K, Pluciennik E, Bednarek AK. WWOX Tumor Suppressor Gene in Breast Cancer, a Historical Perspective and Future Directions. Frontiers in oncology. 2018;8:345.

23. Chang R, Song L, Xu Y, Wu Y, Dai C, Wang X, et al. Loss of Wwox drives metastasis in triple-negative breast cancer by JAK2/STAT3 axis. Nat Commun. 2018;9(1):3486.

24. Inoue K, Fry EA. Haploinsufficient tumor suppressor genes. Adv Med Biol. 2017;118:83–122.

25. Dietlein F, Weghorn D, Taylor-Weiner A, Richters A, Reardon B, Liu D, et al. Identification of cancer driver genes based on nucleotide context. Nat Genet. 2020;52(2):208–18.

26. Gu H, Xu X, Qin P, Wang J. FI-Net: Identification of Cancer Driver Genes by Using Functional Impact Prediction Neural Network. Front Genet. 2020;11:564839.

27. Drost J, Mantovani F, Tocco F, Elkon R, Comel A, Holstege H, et al. BRD7 is a candidate tumour suppressor gene required for p53 function. Nat Cell Biol. 2010;12(4):380–9.

28. Niu W, Luo Y, Zhou Y, Li M, Wu C, Duan Y, et al. BRD7 suppresses invasion and metastasis in breast cancer by negatively regulating YB1-induced epithelial-mesenchymal transition. J Exp Clin Cancer Res. 2020;39(1):30.

29. Fernandez-Majada V, Welz PS, Ermolaeva MA, Schell M, Adam A, Dietlein F, et al. The tumour suppressor CYLD regulates the p53 DNA damage response. Nat Commun. 2016;7:12508.

30. Pseftogas A, Xanthopoulos K, Poutahidis T, Ainali C, Dafou D, Panteris E, et al. The Tumor Suppressor CYLD Inhibits Mammary Epithelial to Mesenchymal Transition by the Coordinated Inhibition of YAP/TAZ and TGF Signaling. Cancers (Basel). 2020;12(8).

31. Wang BJ, Liu DC, Guo QY, Han XW, Bi XM, Wang H, et al. NUDT21 Suppresses Breast Cancer Tumorigenesis Through Regulating CPSF6 Expression. Cancer Manag Res. 2020;12:3069–78.

32. Banerji S, Cibulskis K, Rangel-Escareno C, Brown KK, Carter SL, Frederick AM, et al. Sequence analysis of mutations and translocations across breast cancer subtypes. Nature. 2012;486(7403):405–9.

33. Malik N, Yan H, Moshkovich N, Palangat M, Yang H, Sanchez V, et al. The transcription factor CBFB suppresses breast cancer through orchestrating translation and transcription. Nat Commun. 2019;10(1):2071.

34. Zhu CY, Li CY, Li Y, Zhan YQ, Li YH, Xu CW, et al. Cell growth suppression by thanatos-associated protein 11(THAP11) is mediated by transcriptional downregulation of c-Myc. Cell Death Differ. 2009;16(3):395–405.

35. Evagelou SL, Bebenek O, Specker EJ, Uniacke J. DEAD Box Protein Family Member DDX28 Is a Negative Regulator of Hypoxia-Inducible Factor 2alpha- and Eukaryotic Initiation Factor 4E2-Directed Hypoxic Translation. Mol Cell Biol. 2020;40(6).

36. Lai WL, Hung WY, Ching YP. The tumor suppressor, TAX1BP2, is a novel substrate of ATM kinase. Oncogene. 2014;33(45):5303–9.

37. Johansson P, Jeffery J, Al-Ejeh F, Schulz RB, Callen DF, Kumar R, et al. SCF-FBXO31 E3 ligase targets DNA replication factor Cdt1 for proteolysis in the G2 phase of cell cycle to prevent re-replication. J Biol Chem. 2014;289(26):18514–25.

38. Adhikary A, Chakraborty S, Mazumdar M, Ghosh S, Mukherjee S, Manna A, et al. Inhibition of epithelial to mesenchymal transition by E-cadherin up-regulation via repression of slug transcription and inhibition of E-cadherin degradation: dual role of scaffold/matrix attachment region-binding protein 1 (SMAR1) in breast cancer cells. J Biol Chem. 2014;289(37):25431–44.

39. Malik MZ, Alam MJ, Ishrat R, Agarwal SM, Singh RK. Control of apoptosis by SMAR1. Mol Biosyst. 2017;13(2):350–62.

40. Noll JE, Jeffery J, Al-Ejeh F, Kumar R, Khanna KK, Callen DF, et al. Mutant p53 drives multinucleation and invasion through a process that is suppressed by ANKRD11. Oncogene. 2012;31(23):2836–48.

41. Li J, Belogortseva N, Porter D, Park M. Chmp1A functions as a novel tumor suppressor gene in human embryonic kidney and ductal pancreatic tumor cells. Cell Cycle. 2008;7(18):2886–93.

